# Prion-like transmission of human tau strains in the mouse brain

**DOI:** 10.64898/2026.03.30.715211

**Authors:** Sofia Lövestam, Aki Shimozawa, Airi Tarutani, Reiko Ohtani, Masami Masuda-Suzukake, Kazuko Hasegawa, Andrew C. Robinson, Yuko Saito, Shigeo Murayama, Mari Yoshida, Hisaomi Suzuki, Mitsumoto Onaya, Masato Hasegawa, Michel Goedert, Sjors H.W. Scheres

## Abstract

Most neurodegenerative diseases are believed to spread through the brain by prion-like mechanisms, where filamentous protein assemblies self-propagate by templated seeding ^1^. Distinct conformations of amyloid filaments are thought to provide the physical basis for the strains that lead to different diseases ^2^. However, a central pillar of the prion hypothesis, that strains retain their structural identity upon transmission, has not been demonstrated. Here we show that the injection of tau filaments from the brains of individuals with Alzheimer’s disease or corticobasal degeneration into the brains of wildtype mice leads to the seeded assembly of amyloid filaments made of mouse tau with the same structures as those of the seeds. Thereby, we establish that, like prion strains, tau filaments propagate through templated seeding, and that the mouse is a suitable model to study the molecular mechanisms by which distinct tau folds drive disease-specific pathology in the brain.

---

Misfolded filamentous aggregates of the prion protein cause neurodegeneration in diseases such as Kuru, Creutzfeldt-Jakob disease (CJD), variant CJD, Bovine Spongiform Encephalopathy, Chronic Wasting Disease and Scrapie ^3,4^. Misfolded prion protein assemblies are believed to self-propagate by templated seeding, whereby small amounts of filaments induce the misfolding of the native prion protein into filaments with the same conformation. This ability to self-propagate underlies the prion spread within tissues, between individuals of a given species and, depending on the prion protein sequence similarity, between individuals of distinct species. Various misfolded conformations of the prion protein give rise to prion strains that lead to different incubation times, lesion profiles and clinical phenotypes. Electron cryo-microscopy (cryo-EM) structures of prion filaments have revealed that distinct amyloid folds provide the physical basis for prion strains ^5–9^.

More than twenty neurodegenerative diseases are defined by the filamentous assembly of hyperphosphorylated tau ^10^. Most disease cases with abundant tau filaments are sporadic, with Alzheimer’s disease (AD) being the most common tauopathy. Mutations in *MAPT*, the gene encoding tau, cause dominantly inherited forms of frontotemporal dementias with abundant tau inclusions, consistent with tau filament formation causing neurodegeneration. Cryo-EM structures of tau filaments from human brains have shown that distinct tau folds also characterise different diseases ^11^, and that filaments with a single tau fold accumulate in the brains of individuals with a given tauopathy ^12^. Moreover, small amounts of tau filaments can induce the assembly of normally disordered tau monomers into new filaments ^13,14^. Further support for the prion-like spreading of tau is provided by seeding animal models, in which tau filaments are injected into the mouse brain. Filaments are taken up by brain cells at the injection sites, after which tau pathology spreads to distal regions, suggesting cell-to-cell propagation of tau filaments in the brain ^14–19^. Seeding with human brain-derived tau from different tauopathies recapitulates the cell type-specific characteristics of those diseases ^15^. Subsequent studies showed that these inclusions are robustly recapitulated in the wildtype mouse brain ^17,20,21^ and in mice expressing all six human brain tau isoforms ^22^. Moreover, the injection of AD tau seeds into mice with abundant amyloid-β plaques led to a marked enhancement of tau inclusions compared with injections into wildtype mice ^23^. The injection of filaments assembled from recombinant tau into the brains of wildtype mice and of mice expressing all six human brain tau isoforms showed that filaments with the AD-specific fold seed better than filaments with folds that have not been observed in human brains ^24^.

However, it has so far not been shown that tau folds lead to distinct diseases by prion-like mechanisms and that conformational strains retain their identity upon transmission. Human and mouse sequences are identical within the ordered cores of tau filaments from human brains, indicating that there may be no species barrier for the transmission of human disease-specific tau strains to the mouse. But the atomic structures of tau filaments formed in the seeding mouse models described above remain unknown. To date, cryo-EM structures of tau filaments from transgenic mouse models have differed from those observed for filaments extracted from human brains ^25,26^. Moreover, the intracerebral injection of recombinant wildtype α-synuclein filaments into the brains of mice transgenic for A53T human α-synuclein produced inclusions reminiscent of those in multiple system atrophy (MSA), but the structures of the recombinant filaments were distinct from those observed in MSA, and the seeded filaments only partially retained the structure of the seeds ^27^.

Here we report that the intracerebral injection of wildtype mice with tau filaments extracted from human brains of individuals with AD or CBD results in cell-type specific tau inclusions that are characteristic of both diseases, and that these inclusions comprise filaments of endogenous mouse tau with the same structures as those of the seeds. Our results establish that distinct tau folds act as prion strains in propagating by templated seeding to drive disease-specific pathology in the brain.

## Results

### Injection of AD and CBD tau seeds yields cell type-specific tau pathology in wildtype mice

We injected tau seeds extracted from the frontal cortex of humans with AD or CBD into the striatum of 6-18-week-old wildtype mice (**Figure 1; Methods; Supplementary Tables 1-2**). At 9 months post-injection (mpi) seeded tau pathology was detected using a mouse tau-specific antibody (mTau) and phosphorylation-dependent antibodies AT8 and AT100, indicating the accumulation of phosphorylated mouse tau. A human tau-specific antibody (HT7) gave no staining (**Supplementary Figure 1**). AT8-positive tau pathology had spread from the injection site to the cerebral cortex, corpus callosum and other brain regions for both types of seeds, but the regional distribution and morphology of tau pathology differed depending on the seed. Mice injected with AD seeds showed tau pathology that was confined to neuronal cell bodies and their processes. In contrast, mice injected with CBD seeds showed tau pathology in both nerve cells and glial cells, forming coiled bodies and giving rise to plaque-like astrocytic inclusions. These findings indicate that tau seed structure influences both cellular targeting and pathological morphology *in vivo*.

**Figure 1.**
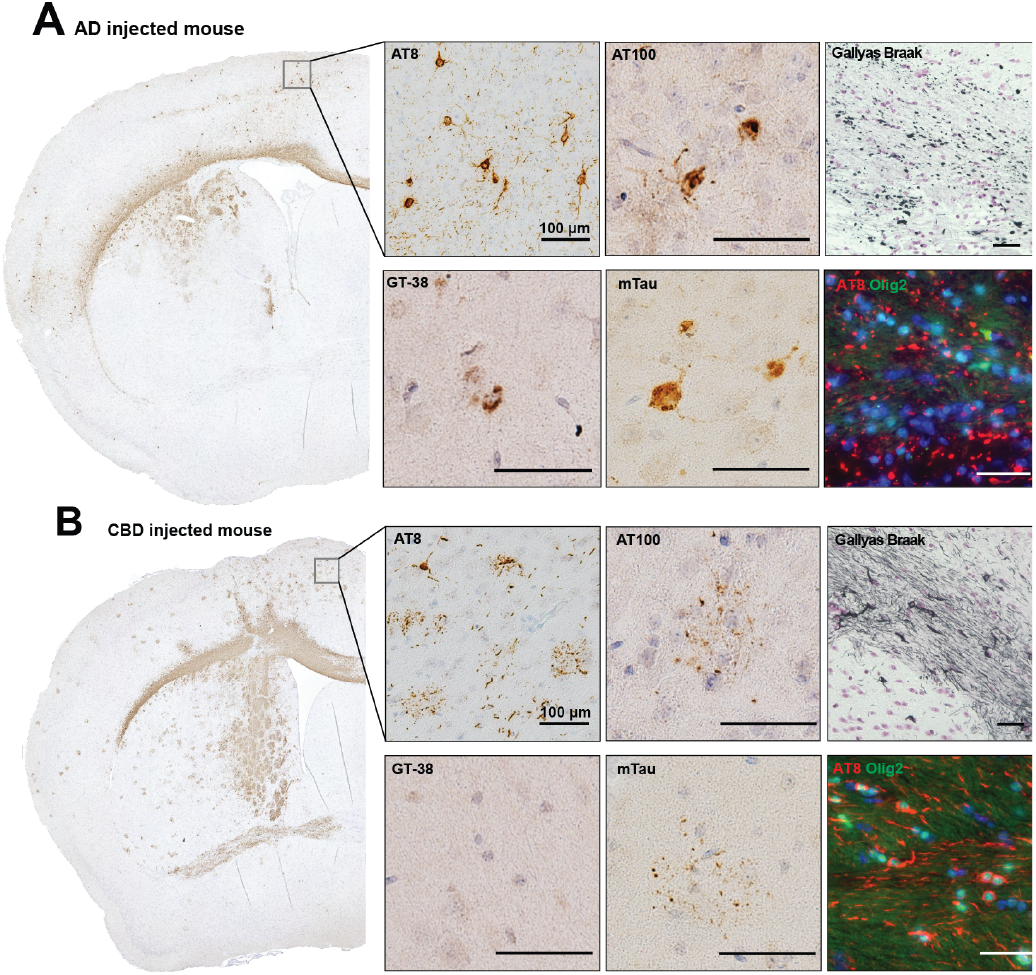
Histological staining of wildtype mouse brains 9 months after injection with human AD or CBD tau seeds. **(A)** AD and **(B)** CBD seed-injected brains stained with antibodies AT8, AT100, mTau and GT-38; stained with Gallyas-Braak; and fluorescence immunostained with AT8 (red) and Olig2 (green). DAPI is in blue. Zoomed-in images show the supplementary motor cortex on the dorsomedial surface of the frontal cortex. Fluorescence staining images show the corpus callosum. Scale bar, 100 *µ*m.

Staining with GT-38, an antibody that recognises tau inclusions in AD but is negative in 3R-only or 4R-only tauopathies ^28^, was positive in mice injected with AD seeds, but negative in those injected with CBD seeds. Immuno-EM with GT-38 was also negative for recombinant filaments with the AD fold made exclusively of 3R tau, but positive for filaments with the CTE fold made exclusively of 4R tau or the AD fold consisting of a mixture of 3R and 4R tau (**Supplementary Figure 2**). These findings indicate that the epitope of the GT-38 antibody lies within the second microtubule-binding repeat (R2), which is absent in 3R tau. Because mice express exclusively 4R tau in the adult brain, these results indicate that the epitope in R2 is accessible for GT-38 binding in the filaments from mouse brains injected with AD tau seeds, but that the same epitope is not accessible in the filaments from mice injected with CBD tau seeds, because it forms part of the ordered filament core.

### Seeded aggregates comprise phosphorylated mouse tau filaments

Western blotting of sarkosyl-insoluble fractions from the injected mouse brains were positive for mTau and negative for HT7, suggesting that the insoluble fractions consisted only of mouse tau, in agreement with the immunohistochemistry described above. Distinct bands with the T46 and AT8 antibodies suggested biochemical differences between insoluble tau seeded with AD or CBD tau. Of note, seeding with CBD tau led to a 37 kDa band that is characteristic of CBD pathology (**Figure 2**). Immuno-EM with mTau and AT8 antibodies of the sarkosyl-insoluble fractions confirmed that these fractions contained filaments composed of phosphorylated mouse tau. Tau filaments from mouse brains injected with CBD seeds had longer cross-over distances than those from brains injected with AD seeds (**Figure 2**).

**Figure 2.**
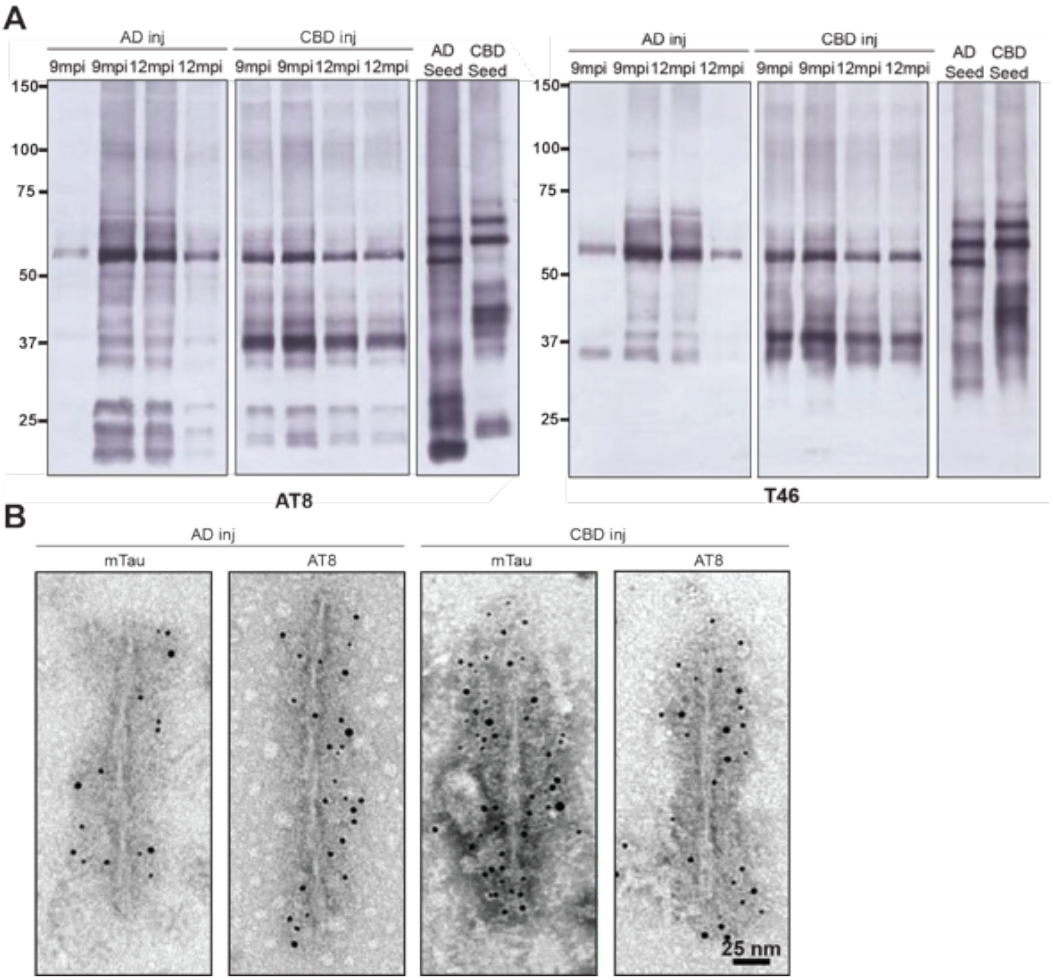
Biochemical characterisation of seeded mouse tau. **(A)** Western blots of sarkosyl-insoluble fractions from brain hemipheres using antibodies AT8 and T46 shows accumulation of insoluble mouse tau at 9 and 12 months post-injection (mpi) for AD and CBD seed-injected WT mice. **(B)** Negative stain electron microscopy with immunogold labelling of AD and CBD-injected mice 12 mpi with antibodies AT8 and mTau. Scale bar, 25 nm.

### Seeded aggregation with CBD tau precedes seeding with AD tau

We next performed a time-course analysis of the injected seeds using histological staining, electron microscopy and Western blotting. The human tau seeds were visible by staining immediately after injection, but they could no longer be detected after one week. Clearance of the injected human seeds was confirmed by negative-stain electron microscopy and Western blotting (**Supplementary Figure 3**).

At 1 mpi, AT8, pFTAA and Gallyas-Braak silver staining showed tau inclusions in mice injected with CBD seeds, but tau inclusions were absent from the brains of mice injected with AD seeds (**Figure 3**). In the latter, tau inclusions became only apparent at 3 mpi. In both the AD and CBD-injected mice, tau inclusions had increased in number by 6 mpi and 9 mpi. The time-dependent increase in tau inclusions was confirmed by Western blotting using anti-Tau C, mTau and AT8 antibodies (**Supplementary Figure 3**).

**Figure 3.**
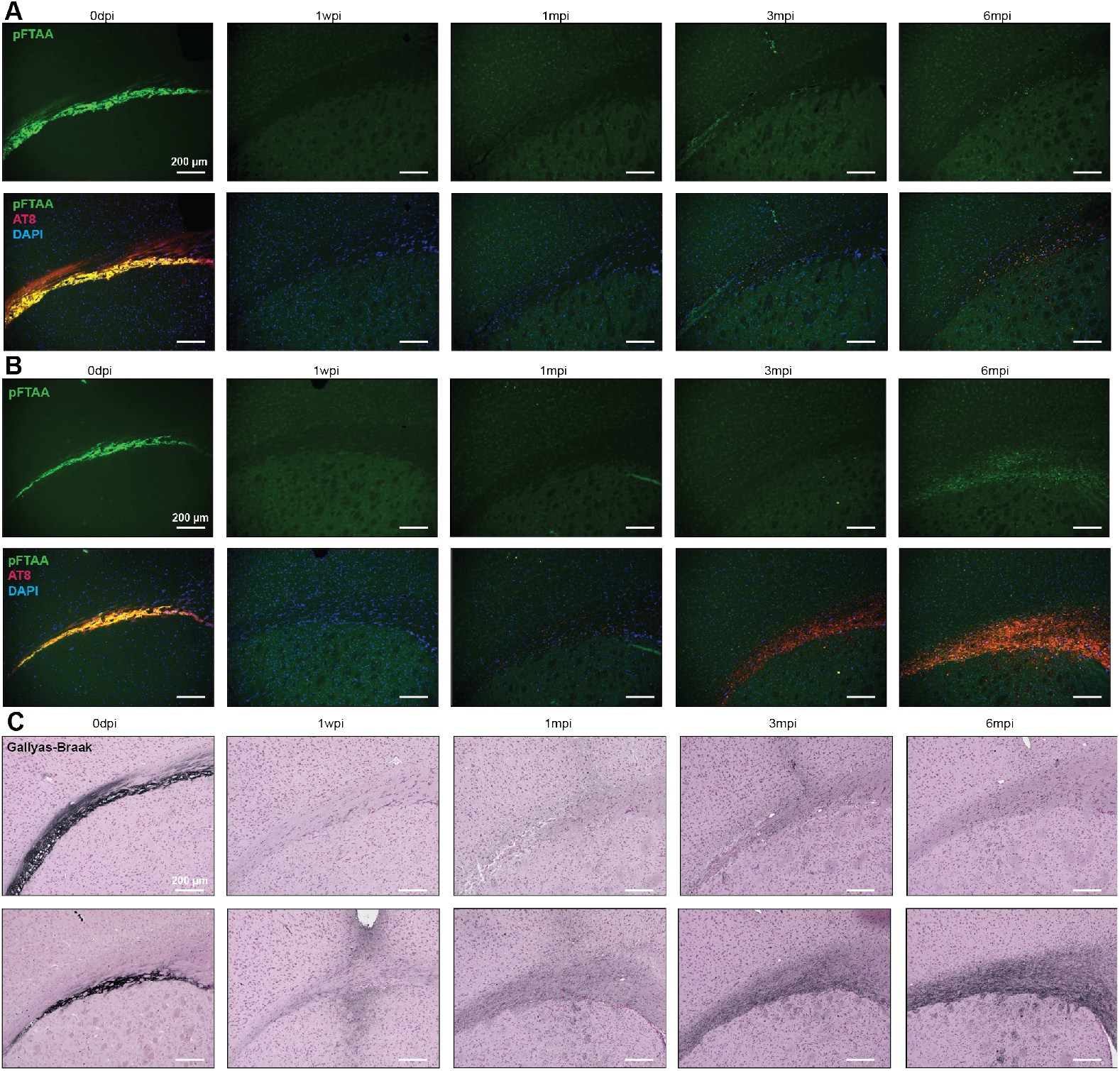
Time-course of tau staining. pFTAA and AT8 staining of the corpus callosum at the level of the frontal cortex from AD **(A)** and CBD **(B)** seed-injected mice immediately after injection (0 dpi), 1 week post-injection (1 wpi), 1month post-injection (1 mpi), 3 mpi and 6 mpi. **(C)** Gallyas-Braak silver staining of AD (top) and CBD (bottom) seed-injected mice at 0 dpi, 1 wpi, 1 mpi, 3 mpi and 6 mpi. Scale bar, 200 *µ*m.

### Seeded tau filaments adopt the same structures as the seeds

Cryo-EM analysis of tau filaments extracted from the brains of mice injected with AD or CBD seeds demonstrated that seeded mouse tau filaments adopted the same structures as those of the human seeds (**Figure 4; Supplementary Figure 4**). In mice injected with AD seeds, the majority of filaments (83%) were paired helical filaments (PHFs), for which we calculated a reconstruction to a resolution of 3.6 Å. This structure was identical to the Alzheimer fold from AD brains ^29,30^. We were unable to obtain a reconstruction for the remaining filaments (17%), which showed 2D class averages that were reminiscent of single protofilaments or of straight filaments (SFs) from AD (**Supplementary Figure 4B**). Cryo-EM images of mouse brains injected with CBD tau seeds contained two filament types: 74% of the filaments were made of a single protofilament and 26% of the filaments were made of two identical protofilaments. We calculated a reconstruction of the singlets to a resolution of 3.4 Å. This structure was identical to human Type I CBD filaments ^31^. A reconstruction of the doublets, to a resolution of 10 Å, suggested that they were identical to type II CBD filaments.

**Figure 4.**
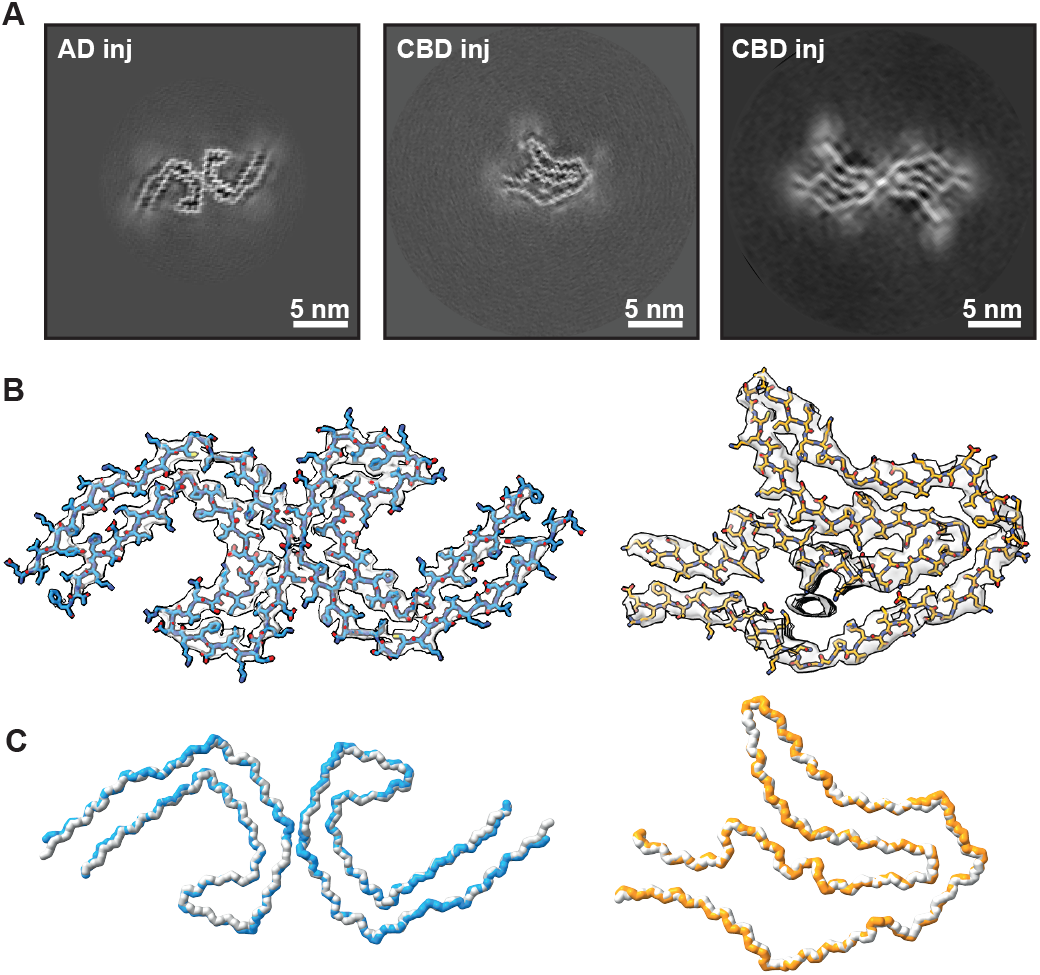
Cryo-EM structures of seeded mouse tau filaments. **(A)** Cryo-EM cross-sections of AD and CBD seed-injected mouse tau filaments perpendicular to the helical axis, each representing one rung. Scale bar, 5 nm. **(B)** The atomic model fitted into the cryo-EM density map (grey transparent) of the AD (blue) and CBD (orange) seed-injected mouse tau filaments. **(C)** Overlay of the C-alpha backbone of AD (blue) and CBD (orange) seed-injected mouse tau filaments overlaid with human *ex vivo* AD PHFs and CBD type I filaments (grey).

## Discussion

We provide evidence for the prion-like transmission of human disease-specific tau strains in the mouse brain. Because the murine and human tau sequences are identical within the ordered cores of the AD and CBD filament folds, mouse tau could assemble into filaments with the same atomic structures as those from human brains. Consistent with previous observations ^15,17,21,22^, and supporting the hypothesis that distinct tau folds represent unique strains, injection of human brain-derived tau seeds led to cell-type specific lesions in the mouse brain that were characteristic of those of AD and CBD.

Alternative splicing of the *MAPT* transcript leads to the expression of six tau isoforms in the adult human brain, with three isoforms comprising three microtubule-binding repeats (3R) and three isoforms comprising four repeats (4R) ^32^. Human tauopathies are classified according to the isoform composition of their filaments: a mixture of all six isoforms assembles in 3R+ 4R tauopathies like AD, only 3R tau assembles in 3R tauopathies like Pick’s disease and only 4R tau assembles in 4R tauopathies such as CBD. Because adult mice only express 4R tau, the absence of 3R tau represents a species barrier for the transmission of tau seeds with the Pick fold. This barrier does not apply to 3R+ 4R and 4R tauopathies. Therefore, tau seeds from other 3R+ 4R and 4R tauopathies, such as chronic traumatic encephalopathy and progressive supranuclear palsy, may also be transmitted in wildtype mice. Alternatively, mice expressing both 3R and 4R mouse tau ^33^, or mice expressing all six human brain tau isoforms in the absence of endogenous mouse tau ^22,34,35^ can be used to transmit seeds from all human tauopathies, and these mice exhibit more extensive tau pathology than wildtype mice ^24^.

Despite the similarities in the mechanisms by which tau and prion strains propagate, important differences exist between the two proteins. Prion diseases are infectious and can be transmitted between individuals, whereas inter-individual transmission of tauopathies has not been reported. Prion diseases typically also progress much faster than tauopathies. Thus, mice injected with prion seeds can develop severe disease ^4^, whereas mice injected with tau seeds appear healthy, despite the presence of tau inclusions in the brain ^14^. It is possible that mice injected with tau seeds do not live long enough to develop symptoms, or that the amount of tau inclusions that forms in these mice is not sufficient to lead to clinical symptoms. Mice that are transgenic for human tau with mutations that cause frontotemporal dementias spontaneously form abundant tau filaments and develop severe neurodegeneration ^36–39^. Moreover, rhesus macaques injected with tau seeds from the brains of individuals with the 4R tauopathy progressive supranuclear palsy developed motor and behavioural impairments six months after seed injection ^40^.

Filament assembly can be separated into three stages. The first stage involves seed uptake, which may involve cell surface receptors ^13,41,42^ and is probably influenced by the size of tau filaments and their surface properties, including the presence or absence of a fuzzy coat. The second stage consists of the recruitment and templated conversion of monomeric tau within the recipient cell ^43^. This step requires compatibility between the seed, the monomeric tau substrate, including its post-translational modifications, and the cell type environment, which may or may not comprise cofactors that are essential for filament assembly. The third and final stage is the release of the newly formed filaments and their uptake by connected cells, resulting in the stereotypical spread of tau pathology through connected brain regions ^44^. Different tau folds may influence one or more of these stages, thereby shaping both the efficiency of propagation and the cell type specificity observed across tauopathies.

Identifying the molecular factors that modulate tau uptake, templated conversion and propagation will be essential for understanding how distinct tau strains transmit specific pathologies in a prion-like manner. Our observation that the intracerebral injection of wildtype mice with tau seeds from human neurodegenerative diseases leads to the assembly of mouse tau filaments that retain the conformation of the human disease-specific seeds indicates that the mouse is a suitable model for investigating these mechanisms. Their elucidation will lead to a better understanding of disease-specific tau strain behaviour and provide new therapeutic avenues.

## Acknowledgements

We thank the individuals and their families for donating brain tissues, T. Darling, I. Clayson and J. Grimmett for help with high-performance computing and the EM facility of the Medical Research Council (MRC) Laboratory of Molecular Biology for help with cryo-EM data acquisition. We also thank Ms. Mayuko Nagata (Tokyo Metropolitan Institute of Medical Science) for technical assistance. This work was supported by the MRC, as part of United Kingdom Research and Innovation (UKRI) [MC_U105184291 to M.G. and MC_UP_A025-1013 to S.H.W.S.]. It was also supported by the Japan Agency for Medical Research and Development (AMED; JP24wm0625120 to A.T. and JP24dk0207074h0001 to M.H., JP24wm0425019, JP24dk0207074h0001, JP24dk0207074s0201 to Y.S., S.M., M.Y. and 24wm0425019h0004 to H.S); the Japan Science and Technology Agency (JST) Core Research for Evolutional Science and Technology (CREST; JPMJCR18H3 to M.H.); and the Japan Society for the Promotion of Science (JSPS) KAKENHI (JP24H00624 to M.H., JP 22H04923 CoBiA to Y.S., S.M.). The work of the Manchester Brain Bank is supported by Alzheimer’s Research UK and the Alzheimer’s Society through the Brains for Dementia Research (BDR) Programme.

## Author contributions

Study concept and design: M.H., M.G., S.H.W.S.; clinicopathological diagnosis: Y.S., S.M., M.Y., M.O.; antibody generation: M.M-S.; injection and immunohistochemistry: A.S., A.T., R.O.; immunoblotting and immunoelectron microscopy: S.L., A.T., M.H.; electron cryo-microscopy: S.L.; interpretation of the data: S.L., A.S., A.T., M.H., M.G., S.H.W.S.; drafting of the manuscript: S.L., A.S., A.T., M.H., M.G., S.H.W.S.; final manuscript: all authors.

## Data availability

Atomic coordinates for the mouse tau filaments from the AD and CBD-seed injected wildtype mouse brains have been deposited at the Protein Data Bank (accession codes 29OS and 29OU); electron microscopy maps have been deposited at Electron Microscopy Data Bank (accession codes 57273 and 57275). All other materials are available upon request. Requests for materials should be addressed to M.H., M.G. and S.H.W.S.

## Additional Information

For the purpose of open access, the MRC Laboratory of Molecular Biology has applied a CC BY public copyright license to any Author Accepted Manuscript version arising.

## Materials and Methods

### Tau seeds from AD and CBD brains

Sarkosyl-insoluble fractions were prepared from the brains of five individuals with neuropathologically confirmed AD and six individuals with CBD (**Supplementary Table 1**). The pellets from ~0.5 g brain samples were washed with 2 mL sterile saline, followed by ultracentrifugation as described ^45^. The pellets were then sonicated in 120 µL sterile saline using a cup horn sonicator (Sonifier® SFX, Branson) at 35% power for 180 s ^46^. These were the AD and CBD seeds that were used for injection. Informed consent was obtained from the individuals who donated their brains for the study. Human brain tissue was used with permission from the ethical review boards of the relevant institutions and hospitals.

**Supplementary Table 1.** AD and CBD cases used in this study.

| **Individual** | **Gender** | **Age at death** |
| --- | --- | --- |
| AD case 1 | Male | 61 |
| AD case 2 | Female | 65 |
| AD case 3 | Male | 56 |
| AD case 4 | Male | 80 |
| AD case 5 | Female | 59 |
| CBD case 1 | Female | 75 |
| CBD case 2 | Female | 79 |
| CBD case 3 | Female | 74 |
| CBD case 4 | Male | 51 |
| CBD case 5 | Male | 73 |
| CBD case 6 | Male | 65 |

**Supplementary Table 2.** Mice used in this study.

| Number of mice | Seeds injected | Age at injection | Injection volume / mode | Time post-injection | Experiments performed |
| --- | --- | --- | --- | --- | --- |
| 9 | AD case 1 | 18 weeks | 3 μL / bilateral | 9 or 12 months | Cryo-EM & immunohistochemistry |
| 5 | AD case 1 | 8 weeks | 3 μL / bilateral | time-course* | Immunoblot & immunohistochemistry |
| 8 | AD case 2 | 6 or 8 weeks | 5 μL / unilateral | 9 months | Immunoblot & immuno-EM |
| 5 | AD case 3 | 6 or 8 weeks | 5 μL / unilateral | 9 months | Immunoblot |
| 6 | AD case 4 | 6 or 8 weeks | 5 μL / unilateral | 9 months | Immunoblot |
| 4 | AD case 5 | 6 or 8 weeks | 5 μL / unilateral | 9 months | Immunoblot |
| 8 | CBD case 1 | 18 weeks | 3 μL / bilateral | 9 or 12 months | Cryo-EM & immunohistochemistry |
| 5 | CBD case 1 | 8 weeks | 3 μL / bilateral | time-course* | Immunoblot & immunohistochemistry |
| 8 | CBD case 2 | 6 or 8 weeks | 5 μL / unilateral | 9 months | Immunoblot |
| 4 | CBD case 3 | 6 or 8 weeks | 5 μL / unilateral | 9 months | Immunoblot |
| 6 | CBD case 4 | 6 or 8 weeks | 5 μL / unilateral | 9 months | Immuno-EM |
| 6 | CBD case 5 | 6 or 8 weeks | 5 μL / unilateral | 9 months | Immunoblot |
| 4 | CBD case 6 | 6 or 8 weeks | 5 μL / unilateral | 9 months | Immunoblot |

\* For the time course experiments, brains were harvested at 0 days, 1 week, 1month, 3 months and 6 months post-injection

### Intracerebral injection of AD and CBD seeds

Wildtype mice (6-18 weeks old) were used. They were housed under standard conditions with *ad libitum* access to food and water. Under isoflurane anaesthesia, C57BL/ 6J male mice (Japan SLC, Inc.) were fixed in a stereotaxic frame and injected with 3 or 5 µL seeds into the striatum bilaterally (coordinates from bregma: ±0.5 mm anterior, ±2.0 mm lateral, and # 3.0 mm depth) using a Hamilton syringe. Following injection, the animals were returned to their cages and analysed 9-12 months later. For cryo-EM, two mouse brains injected with either AD or CBD tau seeds were pooled. All experimental protocols were approved by the Animal Care and Use Committee of the Tokyo Metropolitan Institute of Medical Science.

### Immunohistochemistry

Mice were anaesthetised with mixed anaesthetic (0.75 mg/ kg of medetomidine, 4 mg/ kg of midazolam, and 5 mg/ kg of butorphanol), perfused with saline, and their brains were collected. For histological analysis, brains were fixed with 10% formalin neutral buffer solution (Wako). Fixed brains were then embedded in paraffin and sectioned at 8 µm. The sections were deparaffinised in xylene and rehydrated through a graded ethanol series. For high-sensitivity detection, sections were autoclaved at 121°C for 10 min and treated with formic acid for 5 min, followed by incubation in 3% hydrogen peroxide in phosphate-buffered saline (PBS) for 15 min to inactivate peroxidases. After blocking with 10% calf serum in PBS with 0.1% TritonX-100, sections were incubated overnight at 4°C with primary antibodies. They were then incubated with biotinylated secondary antibodies (1:1000, Vector laboratories) for 2 h. Immunolabeling was visualized using 3,3’-diaminobenzidine (DAB) and an ABC staining kit (Vector Laboratories). Sections were counterstained with haematoxylin. Images were taken using a BX53 microscope equipped with a DP80 camera (Olympus). The following primary anti-tau antibodies were used: phosphorylation-dependent (AT8, mouse monoclonal, 1:1000; Invitrogen; AT100, mouse monoclonal, 1:1000; Invitrogen), a mouse tau-specific antibody (mTau, rabbit polyclonal, 1:1000; Cosmo Bio), a conformational antibody (GT-38, mouse monoclonal, 1:1000; Abcam), a human tau-specific antibody (HT7, mouse monoclonal, 1:2000; Invitrogen) and an antibody specific for the C-terminus of tau (Anti Tau-C, rabbit polyclonal, 1:3000; Cosmo Bio). For double-label immunofluorescence, deparaffinised sections were incubated overnight at 4 °C with primary antibodies (AT8, mouse monoclonal, 1:1000; Invitrogen; Olig2, rabbit monoclonal, 1:500; Abcam). The sections were then washed and incubated with a cocktail of Alexa Fluor 594–conjugated goat anti-mouse IgG and Alexa Fluor 488–conjugated goat anti-rabbit IgG (1:500; Vector Laboratories). Staining with pentameric formyl thiophene acetic acid (pFTAA; Sigma-Aldrich) was performed by incubating the sections with 6µM pFTAA for 1 h at room temperature following AT8 immunohistochemistry. After further washing, the sections were coverslipped with anti-fade mounting medium (Vectashield, Vector Laboratories) and observed using a BZ-X710 microscope (Keyence).

### Modified Gallyas-Braak silver staining

Deparaffinised sections were oxidized in 0.25% potassium permanganate solution for 15 min. After washing, the sections were bleached in 2% oxalic acid solution. The sections were then immersed for 1 min in silver iodide solution (100 mL distilled water, 4 g sodium hydroxide, 10 g potassium iodide, 1.75 mL 1% silver nitrate). After rinsing with 0.5% acetic acid, the sections were incubated in developer solution for 15–20 min. The reactions were stopped with 0.5% acetic acid, followed by washing and incubation in 1% gold chloride solution for 5 min. After washing, the sections were incubated in a fixer solution for 10 min. Sections were counterstained with 0.1% Kernechtrot. Images were taken using a BX53 microscope equipped with a DP80 camera (Olympus).

### Biochemical and ultrastructural analyses of tau filaments

Mouse brains were frozen on dry ice and stored at – 80 °C. Sarkosyl-insoluble pellets were prepared from ~0.5 g tissue, as described ^47^. The pellets were sonicated in 60 µL saline and aliquots used for immuno-EM. Thirty µL of the samples were treated with 30 µL of 2x sample buffer at 100°C for 3 min, and 5~10 µL loaded for immunoblotting. Immunoblotting and immuno-EM were performed using anti-tau antibodies T46 (1:1000), AT8 (1:1000), mTau (1:2000), and human tau-specific antibody HT7 (1:2000), as described ^45^. Sarkosyl-insoluble fractions extracted from the injected mice were dropped onto carbon-coated 300-mesh copper grids (Nissin EM) and dried. The grids were stained with mTau or AT8 (1:50). Secondary antibody was conjugated to 5 nm gold particles (Cytodiagnostics, 1:100). A drop of 2% phosphotungstic acid was added to the stained grids and electron micrographs were recorded with a JEOL JEM-1400 electron microscope.

### Electron cryo-microscopy

Three µL of sarkosyl-insoluble fractions from wildtype mice that had been injected bilaterally with AD or CBD seeds 9-12 months earlier were applied to glow-discharged (Edwards S150B; Edwards High Vacuum International, Crawley, West Sussex, UK) holey carbon grids (Quantifoil Au R1.2/ 1.3, 300 mesh) that were plunge-frozen in liquid ethane using a Vitrobot Mark IV (Thermo Fisher Scientific) at 100% humidity and 4°C. Cryo-EM images were acquired on a Titan Krios electron microscope operated at 300 kV, equipped with a Falcon4i direct electron detector (Thermo Fisher Scientific) and a Selectris-X energy filter. Images were recorded at a dose of 40 electrons per Å^2^ using EPU software (Thermo Fisher Scientific).

### Cryo-EM image processing

Datasets were processed in RELION using amyloid helical reconstruction procedures ^48^. Movie frames were gain - corrected, aligned and dose - weighted using RELION’s own implementation of the motion correction programme ^49^. Contrast transfer function (CTF) parameters were estimated using CTFFIND-4.1 ^50^. Filaments were picked automatically and extracted in boxes of 768×768 pixels, which were downscaled to 128×128 pixels. Reference-free 2D classification followed by bi-hierarchical classification ^51^ was used to separate distinct filament types and select classes suitable for reconstruction. Standard 3D auto-refinements were performed, including optimisation of the helical twist and rise parameters. Bayesian polishing ^52^ and CTF refinement ^53^ were used to improve the resolution of the reconstructions. Final maps were sharpened using standard postprocessing procedures and their resolutions were calculated based on the Fourier shell correlation (FSC) between two independently refined half-maps at a threshold of 0.143 ^54^ (**Supplementary Figure 4**). Atomic models of human brain-derived tau filaments (PDB-IDs 6HRE ^30^ for PHFs and 6TJO ^31^ for CBD Type I filaments) were rigid-body fitted into the maps and refined using ISOLDE ^55^ (**Supplementary Table 3**).

**Supplementary Table 3.**
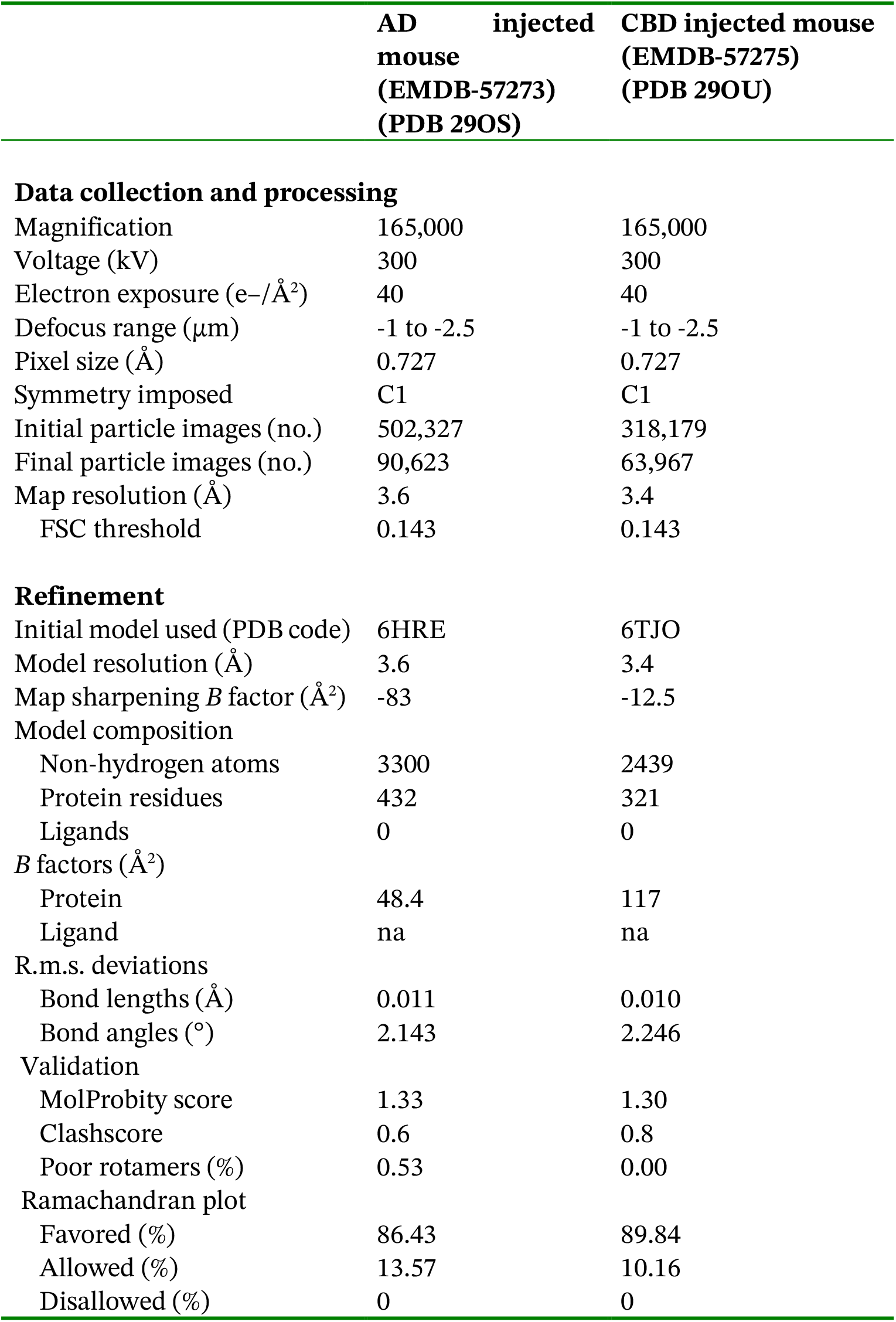
Cryo-EM data collection, refinement and validation statistics.

**Supplementary Figure 1.**
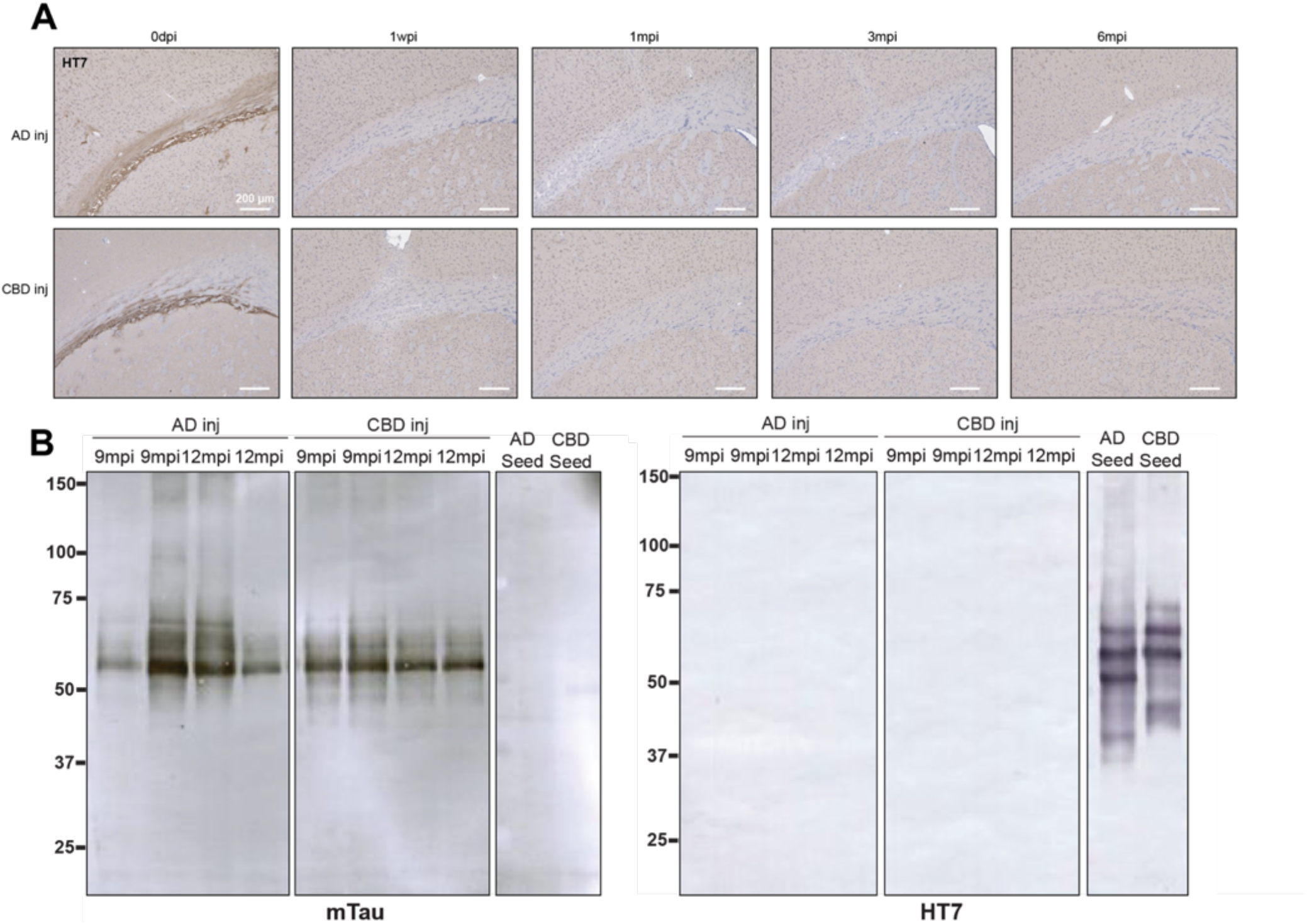
Immunohistochemistry and Western blotting of AD and CBD seed-injected mice 9 and 12 mpi. **A** Immunohistochemistry of the corpus callosum at the level of the frontal cortex with human tau-specific antibody HT7 antibody after intrastriatal injection (0 dpi, 1 wpi, 3 mpi and 6 mpi) of human AD or CBD tau seeds. Scale bar, 200 *µ*m.**B**. Western blotting of sarkosyl-insoluble fractions from brain hemispheres of AD and CBD seed-injected mice at 9 mpi and 12 mpi and of the AD and CBD seeds that were used for injection with antibodies specific for mouse (mTau) or human (HT7) tau.

**Supplementary Figure 2.**
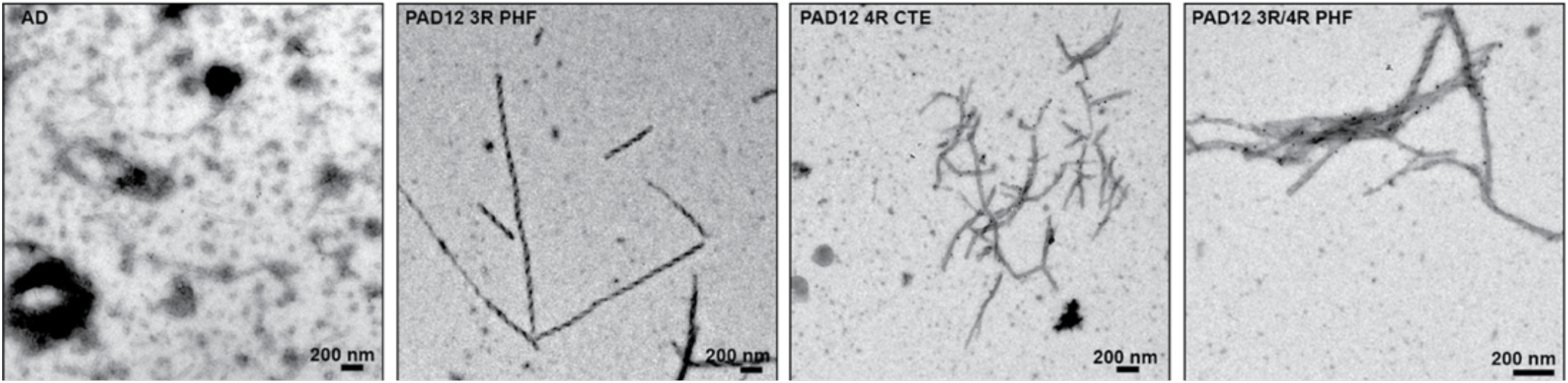
Negative-stain electron microscopy of tau filaments immunogold-labelled with the GT-38 antibody. From left to right: AD-derived filaments, recombinant PAD12 PHFs composed of 3R-only tau, recombinant PAD12 CTE filaments composed of 4R-only tau, and recombinant PAD12 PHFs composed of a mixture of 3R and 4R tau. Scale bar, 200 nm.

**Supplementary Figure 3.**
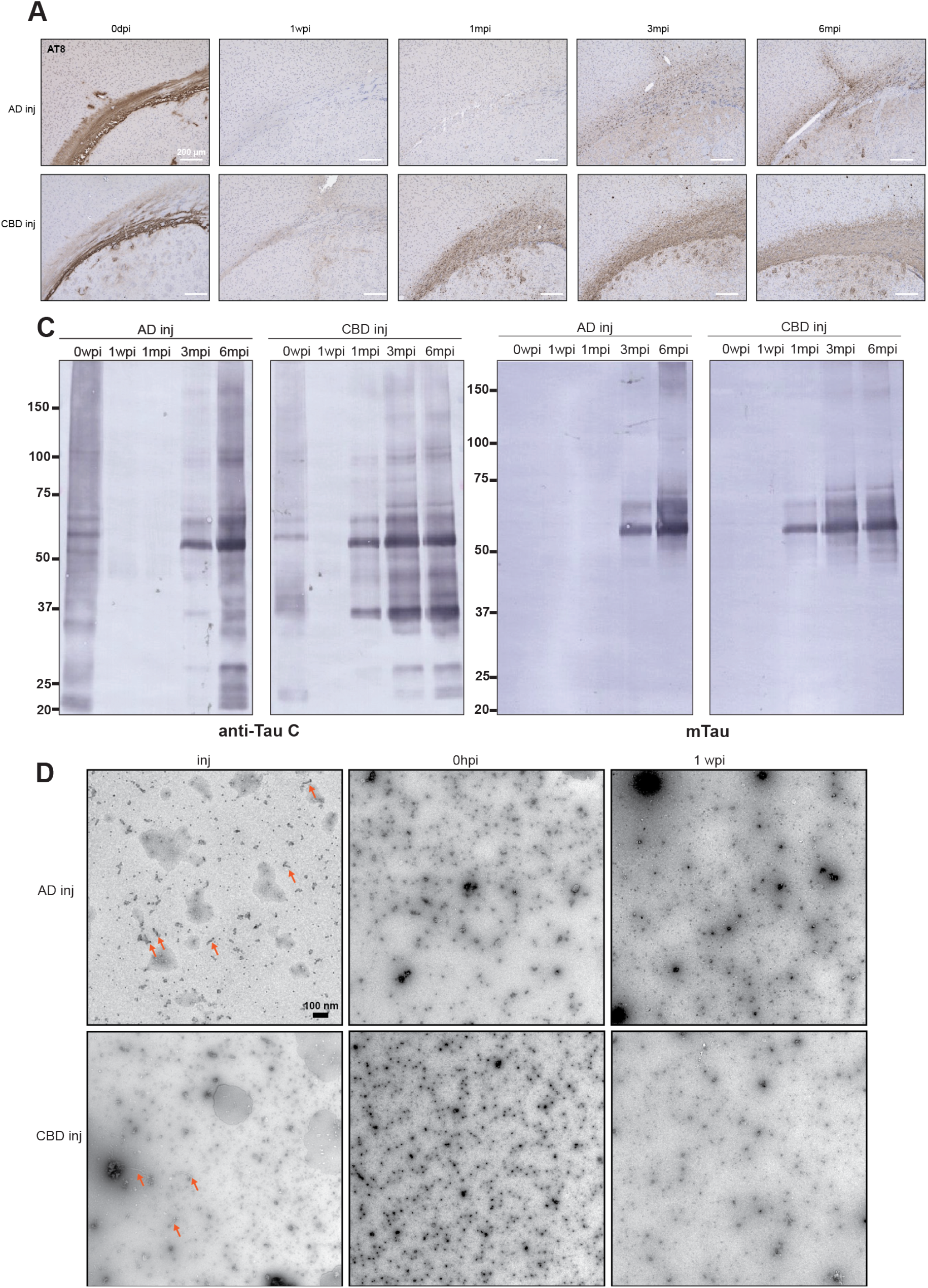
Time course immunohistochemistry, Western blotting and negative stain electron microscopy of AD and CBD seed-injected mice. **A** Immunohistochemistry of the corpus callosum at the level of the frontal cortex with anti-tau antibody AT8 of AD and CBD seed-injected mice 0dpi, 1 wpi, 3 mpi and 6 mpi after intrastriatal injection. Scale bar, 200 µm. **C** Western blotting of sarkosyl-insoluble fractions from brain hemispheres of AD and CBD-injected mice at 0, 1 wpi, 3 mpi and 6 mpi using antibodies anti-Tau C and mTau. **D** Negative stain electron microscopy of the AD and CBD seeds used for injections (inj), immediately after injection (0hpi) and 1 wpi. Arrows point to filaments. Scale bar, 100 nm.

**Supplementary Figure 4.**
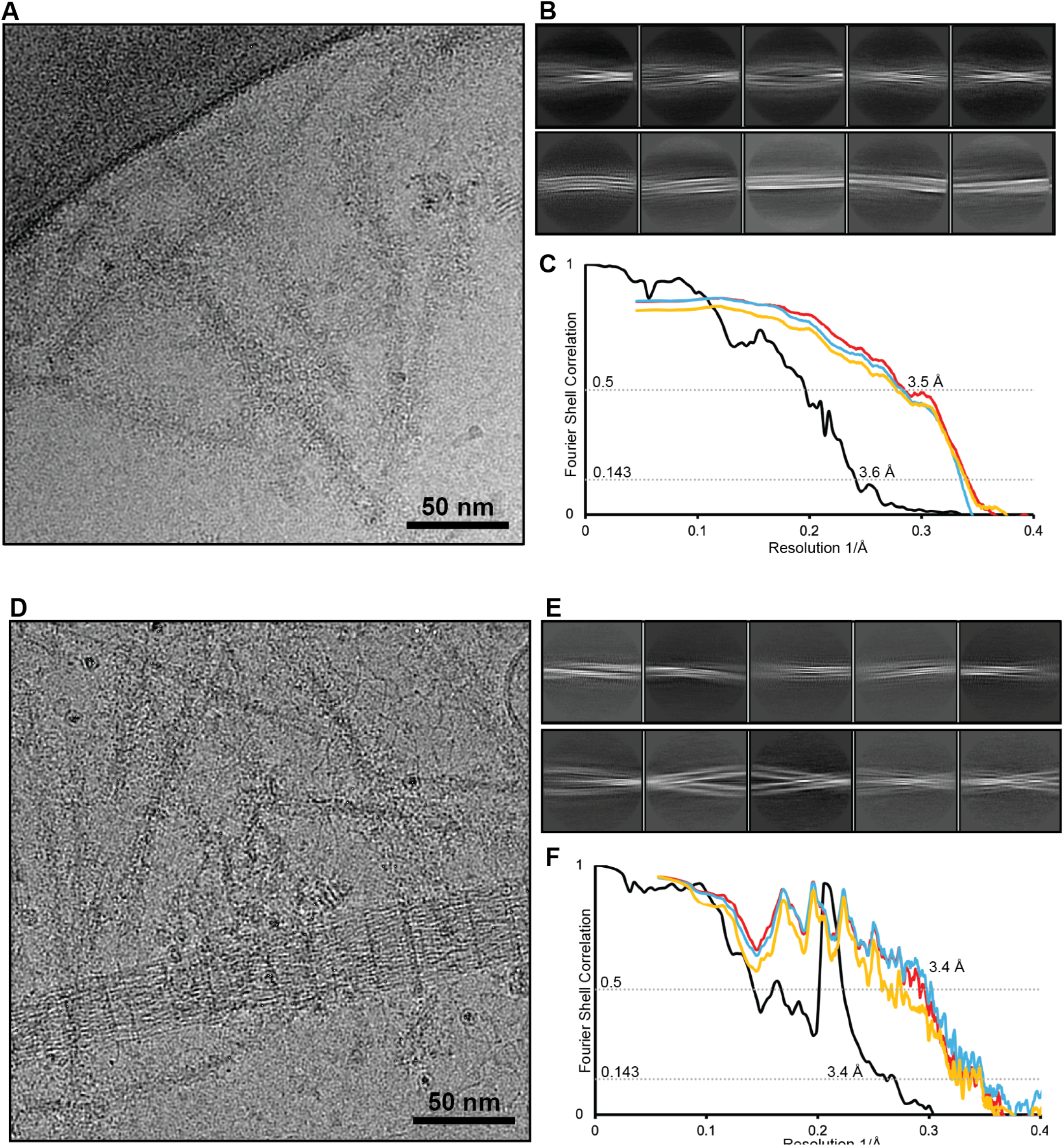
Cryo-EM data processing of seeded mouse tau filaments. **A-C** For AD and **D-F** for CBD seed-injected mice. **A,D** Representative cryo-EM micrographs. Scale bar, 50 nm. **B, E** Representative 2D class averages; the box sizes are 630 Å. **C, F** Fourier shell correlation curves (FSC) between two independently refined half-maps (black); between the atomic model fitted in half-map 1 and half-map1 (blue); between the atomic model fitted in half-map 1 and halfmap 2 (yellow) and between the atomic model fitted in the sum of the two half-maps and the sum of the two half-maps (red).

## References

1. Peng, C., Trojanowski, J. Q. & Lee, V. M.-Y. Protein transmission in neurodegenerative disease. Nat Rev Neurol 16, 199–212 (2020).

2. Collinge, J. & Clarke, A. R. A general model of prion strains and their pathogenicity. Science 318, 930–936 (2007).

3. Benestad, S. L. & Telling, G. C. Chronic wasting disease: an evolving prion disease of cervids. in Handbook of Clinical Neurology vol. 153 135–151 (Elsevier, 2018).

4. Prusiner, S. B. Prions. Proc Natl Acad Sci U S A 95, 13363–13383 (1998).

5. Alam, P. et al. Cryo-EM structure of a natural prion: chronic wasting disease fibrils from deer. Acta Neuropathol 148, 56 (2024).

6. Hoyt, F. et al. Cryo-EM structure of anchorless RML prion reveals variations in shared motifs between distinct strains. Nat Commun 13, 4005 (2022).

7. Kraus, A. et al. High-resolution structure and strain comparison of infectious mammalian prions. Mol Cell 81, 4540-4551.e6 (2021).

8. Manka, S. W. et al. A structural basis for prion strain diversity. Nat Chem Biol 19, 607–613 (2023).

9. Manka, S. W. et al. 2.7 Å cryo-EM structure of ex vivo RML prion fibrils. Nat Commun 13, 4004 (2022).

10. Goedert, M., Crowther, R. A., Scheres, S. H. W. & Spillantini, M. G. Tau and neurodegeneration. Cytoskeleton (Hoboken) 81, 95–102 (2024).

11. Shi, Y. et al. Structure-based classification of tauopathies. Nature 598, 359–363 (2021).

12. Scheres, S. H. W., Ryskeldi-Falcon, B. & Goedert, M. Molecular pathology of neurodegenerative diseases by cryo-EM of amyloids. Nature 621, 701–710 (2023).

13. Frost, B., Jacks, R. L. & Diamond, M. I. Propagation of tau misfolding from the outside to the inside of a cell. J. Biol. Chem. 284, 12845–12852 (2009).

14. Clavaguera, F. et al. Transmission and spreading of tauopathy in transgenic mouse brain. Nat. Cell Biol. 11, 909–913 (2009).

15. Clavaguera, F. et al. Brain homogenates from human tauopathies induce tau inclusions in mouse brain. Proc. Natl. Acad. Sci. U.S.A. 110, 9535–9540 (2013).

16. Boluda, S. et al. Differential induction and spread of tau pathology in young PS19 tau transgenic mice following intracerebral injections of pathological tau from Alzheimer’s disease or corticobasal degeneration brains. Acta Neuropathol 129, 221–237 (2015).

17. Guo, J. L. et al. Unique pathological tau conformers from Alzheimer’s brains transmit tau pathology in nontransgenic mice. J. Exp. Med. 213, 2635–2654 (2016).

18. Iba, M. et al. Synthetic tau fibrils mediate transmission of neurofibrillary tangles in a transgenic mouse model of Alzheimer’s-like tauopathy. J. Neurosci. 33, 1024–1037 (2013).

19. Iba, M. et al. Tau pathology spread in PS19 tau transgenic mice following locus coeruleus (LC) injections of synthetic tau fibrils is determined by the LC’s afferent and efferent connections. Acta Neuropathol 130, 349–362 (2015).

20. Narasimhan, S. et al. Human tau pathology transmits glial tau aggregates in the absence of neuronal tau. J Exp Med 217, e20190783 (2020).

21. Narasimhan, S. et al. Pathological tau strains from human brains recapitulate the diversity of tauopathies in nontransgenic mouse brain. J. Neurosci. 37, 11406–11423 (2017).

22. He, Z. et al. Transmission of tauopathy strains is independent of their isoform composition. Nat Commun 11, 7 (2020).

23. He, Z. et al. Amyloid-! plaques enhance Alzheimer’s brain tau-seeded pathologies by facilitating neuritic plaque tau aggregation. Nat Med 24, 29–38 (2018).

24. Kasen, A. et al. Seed structure and phosphorylation in the fuzzy coat impact tau seeding competency. Nat Commun 16, 9240 (2025).

25. Schweighauser, M. et al. Cryo-EM structures of tau filaments from the brains of mice transgenic for human mutant P301S Tau. Acta Neuropathol Commun 11, 160 (2023).

26. Zhao, W. et al. Cryo-EM structures reveal variant Tau amyloid fibrils between the rTg4510 mouse model and sporadic human tauopathies. Cell Discov 10, 27 (2024).

27. Burger, D. et al. Synthetic α-synuclein fibrils replicate in mice causing MSA-like pathology. Nature 648, 409–417 (2025).

28. Gibbons, G. S. et al. Detection of Alzheimer disease (AD)-specific tau pathology in AD and NonAD tauopathies by immunohistochemistry with novel conformation-selective tau antibodies. J Neuropathol Exp Neurol 77, 216–228 (2018).

29. Fitzpatrick, A. W. P. et al. Cryo-EM structures of tau filaments from Alzheimer’s disease. Nature 547, 185–190 (2017).

30. Falcon, B. et al. Tau filaments from multiple cases of sporadic and inherited Alzheimer’s disease adopt a common fold. Acta Neuropathol. 136, 699–708 (2018).

31. Zhang, W. et al. Novel tau filament fold in corticobasal degeneration. Nature 580, 283–287 (2020).

32. Goedert, M., Spillantini, M. G., Jakes, R., Rutherford, D. & Crowther, R. A. Multiple isoforms of human microtubule-associated protein tau: sequences and localization in neurofibrillary tangles of Alzheimer’s disease. Neuron 3, 519–526 (1989).

33. Hosokawa, M. et al. Development of a novel tau propagation mouse model endogenously expressing 3 and 4 repeat tau isoforms. Brain 145, 349–361 (2022).

34. Andorfer, C. et al. Hyperphosphorylation and aggregation of tau in mice expressing normal human tau isoforms. J Neurochem 86, 582– 590 (2003).

35. Saito, T. et al. Humanization of the entire murine Mapt gene provides a murine model of pathological human tau propagation. J Biol Chem 294, 12754–12765 (2019).

36. Allen, B. et al. Abundant tau filaments and nonapoptotic neurodegeneration in transgenic mice expressing human P301S tau protein. J Neurosci 22, 9340–9351 (2002).

37. Götz, J., Chen, F., Barmettler, R. & Nitsch, R. M. Tau filament formation in transgenic mice expressing P301L tau. J Biol Chem 276, 529–534 (2001).

38. Lewis, J. et al. Neurofibrillary tangles, amyotrophy and progressive motor disturbance in mice expressing mutant (P301L) tau protein. Nat Genet 25, 402–405 (2000).

39. Yoshiyama, Y. et al. Synapse loss and microglial activation precede tangles in a P301S tauopathy mouse model. Neuron 53, 337–351 (2007).

40. Darricau, M. et al. Tau seeds from patients induce progressive supranuclear palsy pathology and symptoms in primates. Brain 146, 2524– 2534 (2023).

41. Guo, J. L. & Lee, V. M.-Y. Seeding of normal tau by pathological tau conformers drives pathogenesis of Alzheimer-like tangles. J Biol Chem 286, 15317–15331 (2011).

42. Rauch, J. N. et al. LRP1 is a master regulator of tau uptake and spread. Nature 580, 381–385 (2020).

43. Sanders, D. W. et al. Distinct tau prion strains propagate in cells and mice and define different tauopathies. Neuron 82, 1271–1288 (2014).

44. Braak, H. & Braak, E. Neuropathological stageing of Alzheimer-related changes. Acta Neuropathol. 82, 239–259 (1991).

45. Tarutani, A. et al. Human tauopathy-derived tau strains determine the substrates recruited for templated amplification. Brain 144, 2333– 2348 (2021).

46. Tarutani, A., Arai, T., Murayama, S.Hisanaga, S.-I. & Hasegawa, M. Potent prion-like behaviors of pathogenic α-synuclein and evaluation of inactivation methods. Acta Neuropathol Commun 6, 29 (2018).

47. Taniguchi-Watanabe, S. et al. Biochemical classification of tauopathies by immunoblot, protein sequence and mass spectrometric analyses of sarkosyl-insoluble and trypsin-resistant tau. Acta Neuropathol. 131, 267–280 (2016).

48. He, S. & Scheres, S. H. W. Helical reconstruction in RELION. J. Struct. Biol. 198, 163–176 (2017).

49. Zivanov, J. et al. New tools for automated high-resolution cryo-EM structure determination in RELION-3. Elife 7, (2018).

50. Rohou, A. & Grigorieff, N. CTFFIND4: Fast and accurate defocus estimation from electron micrographs. J Struct Biol 192, 216–221 (2015).

51. Lövestam, S., Shi, J., Li, D., Jamali, K. & Scheres, S. H. W. Cryo-EM image processing of amyloid filaments in RELION-5.1. Preprint at 10.64898/2026.03.17.712386 (2026). Is this in BioRxiv?

52. Zivanov, J., Nakane, T. & Scheres, S. H. W. A Bayesian approach to beam-induced motion correction in cryo-EM single-particle analysis. IUCrJ 6, 5–17 (2019).

53. Zivanov, J., Nakane, T. & Scheres, S. H. W. Estimation of high-order aberrations and anisotropic magnification from cryo-EM data sets in RELION-3.1. IUCrJ 7, 253–267 (2020).

54. Scheres, S. H. W. & Chen, S. Prevention of overfitting in cryo-EM structure determination. Nat Meth 9, 853–854 (2012).

55. Croll, T.I. ISOLDE: a physically realistic environment for model building into low-resolution electron-density maps. Acta Crystallogr D Struct Biol 74, 519–530 (2018).

